# Strain dynamics of specific contaminant bacteria modulate the performance of ethanol biorefineries

**DOI:** 10.1101/2021.02.07.430133

**Authors:** Felipe Senne de Oliveira Lino, Maria-Anna Misiakou, Kang Kang, Simone S. Li, Bruno Labate Vale da Costa, Thiago Olitta Basso, Gianni Panagiotou, Morten Otto Alexander Sommer

## Abstract

Bioethanol is a viable alternative for fossil fuels, and its use has lowered CO_2_ emissions by over 500 million tonnes in Brazil alone by replacing more than 40% of the national gasoline consumption. However, contaminant bacteria reduce yields during fermentation. Our understanding of these contaminants is limited to targeted studies, and the interplay of the microbial community and its impact on fermentation efficiency remains poorly understood. Comprehensive surveying and longitudinal analysis using shotgun metagenomics of two major biorefineries over a production season revealed similar patterns in microbial community structure and dynamics throughout the entire fermentation system. Strain resolution metagenomics identified specific *Lactobacillus fermentum* strains as strongly associated with poor industrial performance and laboratory-scale fermentations revealed yield reductions of up to 4.63±1.35% depending on the specific contaminating strains. Selective removal of these strains could reduce emissions from the bioethanol industry by more than 2×10^6^ tonnes per year. Using the large-scale Brazilian ethanol fermentations as a model system for studying microbiome-phenotype relationships this study further demonstrates how high-resolution metagenomics can identify culprits of large scale industrial biomanufacturing.

## Introduction

The Brazilian sugarcane ethanol production process generates more than 30 billion liters of ethanol per year. This corresponds to more than 40% of all the energy consumed by light vehicles in Brazil replacing the demand for almost 550 million barrels of oil per year^1^. The use of this biofuel, either as a sole fuel or blended in the gasoline, results in reductions of more than 60% in total greenhouse gases emissions^1,2^. The sugarcane ethanol production process deploys specific *Saccharomyces cerevisiae* strains, in very high cell density fed-batch fermentations operated with cell recycling^3,4^. Usually the production season lasts for 250 consecutive days per year, with an average mill performing up to 3 cycles of fermentation per day^4^. In total yeast cell populations in a single fermentation exceed 10^8^ cells x ml^-1 4^ Still, contamination of this non-sterile process remains a major problem leading to overall yield reductions of 3%, corresponding to over 960 million liters of ethanol ^5^.

Contamination is mainly caused by lactic acid bacteria already present on the raw material which tolerate ethanol, low pH and high temperature^6^. To control the bacterial contamination, yeast cells are acid washed after every fermentation cycle before re-innoculation^4^. In spite of such measures contamination continues to compromise the industrial process^4,5,7^. To further address this issue, antibiotics and other antimicrobial compounds are used for contamination control. However, antibiotic use raises serious concerns with regards to the global antibiotic resistance crisis and also negatively impacts process economics. Given the continuous problems with contamination there is an increasing need to understand the contaminating microbial community in these bioprocesses as well as its effect on yeast fermentations^3–5,8^.

Microbial communities are integral parts of most natural processes, from biogeochemical cycles to the human health ^9,10^, and the interactions among populations within these communities often shape their functionalities and the surrounding environment^11^. To date, industrial fermentation microbiomes have been studied with limited resolution applying either culture based methods^12–16^ or culture independent methods, like metabarcoding, focusing mainly in a specific process steps^17,18^. These studies have either tried mainly to understand the overall composition of the microbial community^13,17,18^, or to understand the impact of specific contaminant species in ethanol fermentations at controlled laboratory environments^6,19,20^. Yet, new studies are needed to correlate the composition of a complex community to actual industrial process performance, and to discern the potential impact of strain-level variations in the functionalities of such contaminating microbiomes^21^. Shotgun metagenomic sequencing could be a valuable tool for pinpointing the contaminants that most significantly affect the performance of currently established bioprocesses^22^.

In the present work, we have sampled all the unitary steps of the ethanol production process of two mills in Brazil, during an entire fermentation season. We use shotgun metagenomics and cultivation-based approaches to analyze the microbial community composition and pinpoint specific detrimental strains configurations negatively impacting overall process performance, as well as the mechanisms governing the community dynamics. This set of new information reveals that higher-resolution metagenomics analysis are critical for understanding the dynamics of microbial communities, and that strain level modifications are responsible for perturbing a stablished microbiome.

## Results

### Independent sugarcane biorefineries share similar microbiome dynamics

We selected two independently operated sugarcane ethanol mills in Brazil, hereafter referred to as Mill A and Mill B, located over 300 km from each other but situated in similar climate regions (**Methods**). The mills are also similar in overall production capacity and deploy the Melle-Boinot fermentation process (**Figure 1A**)^4^. In this process ethanol is produced via fast, high cell-density, fed-batch fermentations. After the fermentation is finished yeast biomass is recovered via centrifugation. This yeast cream is transferred to a separate vat, diluted with water and acid washed to kill the contaminant bacteria. After this treatment, the yeast cream is pumped back to the fermenters, and the process starts over for as many as 750 fermentation batches per year^4^. The fermentation process is comprised of unidirectional steps providing defined sampling points for our analysis (**Figure 1A**). To reduce potential bias introduced by seasonal variation, we sampled each mill at three distinct timepoints throughout the production season (**Supplementary Table 1**). We also collected fermentation metrics relevant to evaluate the ethanol production process performance (**Supplementary Table 1**). In total, shotgun metagenomic sequencing was applied to 56 samples yielding more than 2.8×10^5^ Gbp high quality data (**Supplementary Table 2, Methods**).

**Figure 1:**
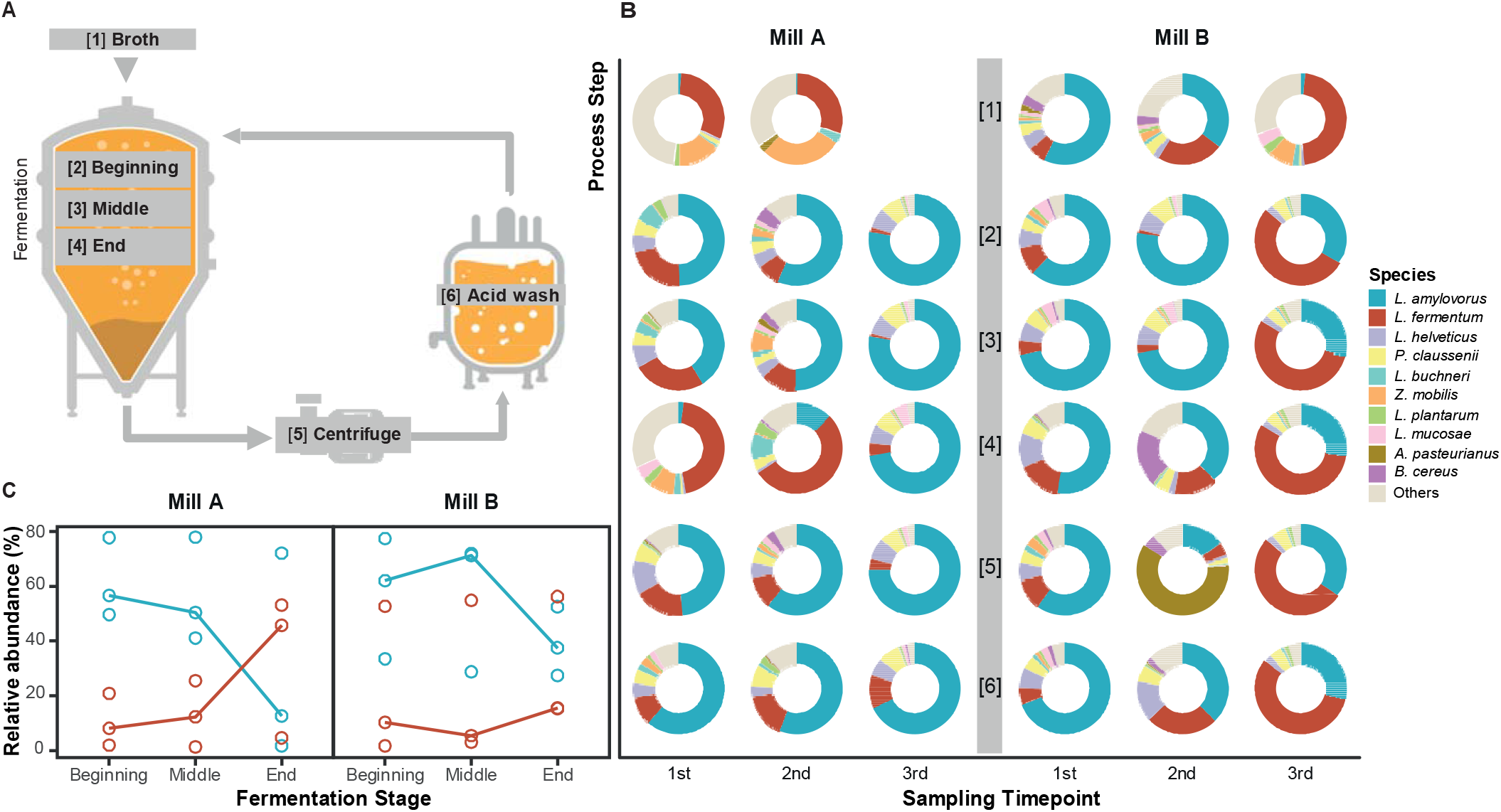
Sampling strategy and competition between *L. amylovorus* and *L. fermentum* in the bioethanol fermentation process. **A:** Schematic of the fermentation process, indicating sampled steps: 1. Broth: Feeding line with fresh fermentation media; 2. Fermentation beginning: the beginning until middle of vessel feeding time (e.g. time 0 to 1.5h of feeding); 3. Middle: the middle until the end of feeding (e.g. time 1.5 to 3h of feeding); 4: End: the final hours of fermentation, after feeding ceased; 5. Centrifuge: the yeast cream, resulting from the separation of the wine sent to distillation; 6. Acid wash: samples collected by the end of this treatment. Vector images were obtained from Flaticon (www.flaticon.com). **B:** Species-level microbial community showing the 10 most prevalent contaminants across sampling timepoints (x-axis) and process steps (y-axis) in Mills A (left) and B (right), expressed as relative abundances. Beige color indicates all remaining species of the community. **C:** Relative abundances of *L. amylovorus* (blue) and *L. fermentum* (red) across the fermentation steps, as shown in **A**. Spearman’s correlation analysis of their relative abundances suggests competition among these two species (r = −0.908; FDR < 4.3 x 10^-14^). Due to low biomass, DNA extraction was not possible for sample [1] for the 3^rd^ sampling timepoint from Mill A. During the 3^rd^ sampling timepoint, Mill B was operating below its maximum capacity. Biomass was left idle for longer periods in the vessels. This might explain why all the process steps are similar in community composition.

Focusing on the prokaryotic component of the metagenomes (29.16±25.19%), we found Firmicutes to be the most prevalent phylum, owing mostly to a high abundance of *Lactobacillaceae* species (**Figure 1B, Supplementary Table 3, Supplementary Table 4)**^6,13,18,19^. Microbial communities from fermentation broth were found to be the most diverse and least similar to those in the rest of the fermentation process (**Supplementary Figure 1**). This difference is mainly driven by the overall dominance of lactobacilli during the fermentation process, since these organisms are best fit to handle the low oxygen and pH, and high temperature and ethanol concentrations, found in industrial fermentations^19^. In addition, we observed that the composition of the contaminant microbiomes across all industrial process steps were highly similar by the end of the production season (final sampling timepoint). Inspection of the collected data revealed that both mills were operating below maximum capacity due to lack of raw material, which resulted in microbial biomass being left idle for longer periods in the fermentation vessels. Irrespective of this the microbial communities of both mills were not found to differ significantly when comparing across process steps and timepoints (*p*=0.293, PERMANOVA; **Supplementary Figure 2**).

The majority of the microbial communities, throughout the entire fermentation process, were found to be dominated by either *L. fermentum* or *L. amylovorus* (**Figure 1B**). These two species have independently been described as contaminants in other ethanol fermentation processes^6,23,24^. We find that these two species constitute more than 50% of the relative abundance of these contaminating communities, demonstrating how uneven such communities are. Interestingly, when comparing relative abundances, we detected an inverse relationship between the two species during the fermentation stages, which suggests competition between *L. fermentum* and *L. amylovorus* during the ethanol production process (Spearman’s correlation ρ = −0.908; FDR < 4.3 x 10^-14^, **Figure 1C**). For both mills, a similar pattern was observed, where *L. amylovorus* dominates at the beginning of the fermentation followed by a decline in its relative abundance in the community throughout the fermentation process.

In contrast *L. fermentum* expands its relative abundance in the community from the beginning to the end of the fermentation. The tolerance of *L. fermentum* towards high ethanol titres^13^ might partly explain its higher relative abundance in the community during the final stages of fermentation. The heterofermentative metabolism of *L. fermentum* might also be better suited to compete with yeast for nutrients in this fermentation setup, compared to the homofermentative metabolism of *L. amylovorus*^6^.

### Microbial community composition affects industrial process performance

To establish if the dynamics observed within the contaminant microbial community were associated with environmental factors or overall fermentation yield, we incorporated fermentation data and industrial performance indicators into our analyses (**Figure 2A, Supplementary Table 1**). Throughout the bioethanol production season, increased ethanol yield was found to strongly associate with lower acidity titres in the fermentation (Spearman’s correlation ρ = −0.84, FDR = 2.09 x 10^-5^), while increases in bacteria negatively impacted the yeast viability (ρ = −0.72, FDR = 2 x 10^-3^).

**Figure 2:**
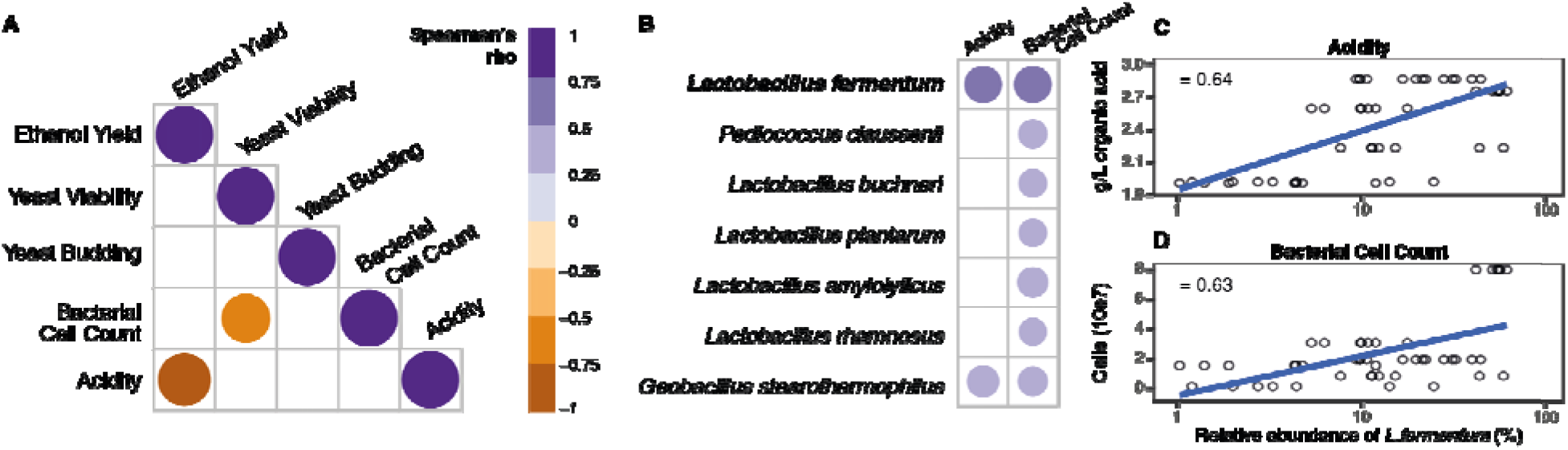
Microbial species and other factors that influence industrial fermentation performance. **A:** Fermentation parameters that showed strong associations throughout the production season. Positive correlations are depicted as purple, whereas negative correlations are depicted as orange. Size of each point denotes the strength of correlation. Increased acidity is linked with lower ethanol yield, and increased number of bacterial cells reduces yeast viability. B: Microbial species associated with fermentation performance. Increased *L. fermentum* during fermentation in strongly linked to higher acidity titres. Species are ordered by decreasing relative abundance. **C:** Correlation between relative abundance of *L. fermentum* and acidity (ρ=0.64; FDR = 1.5 x 10^-5^). **D:** Correlation between relative abundance of *L. fermentum* and bacterial cell count (ρ= 0.63; FDR = 9.5 x 10^-6^).

Based on these findings, we then sought to identify contaminant species that were associated with fermentation performance (**Methods**). We did not expect to find strong associations between the species abundance and ethanol yield, due to well-documented caveats of ethanol quantification methods^25^. Yet, *L. fermentum* was found to be strongly associated with increased acidity titres, which hamper ethanol yield (**Figure 2A and 2B**, Spearman’s correlation ρ = 0.72, FDR < 1.50 × 10^-6^). We also identified a number of bacterial species that were not previously associated with impacts over industrial ethanol production, like *Geobacillus stearothermophillus*. This species is a thermophile capable of producing a vast array of cellulolytic enzymes^26^. Its presence in the fermentation suggests that this industrial environment is a potential untapped source for novel industrially relevant enzymes.

### The fate of fermentations is defined by the interplay of different *L. fermentum* strains during the acid wash

Given its strong association with increased acidity, which creates detrimental fermentation conditions^19^, we sought to establish if the negative effect conferred by *L. fermentum* was mediated by strain variants of this bacteria. SNV profiling of core *L. fermentum* genes revealed the presence of 3 distinct strain families within the species (**Methods, Supplementary Figure 3, Supplementary Table 5**). Applying these profiles to our samples enabled insights into the strain population structure and dynamics during industrial fermentation and its impact on performance.

To more accurately evaluate and compare the performance amongst different fermentation batches^25^, we devised a composite parameter that incorporated ethanol yield measurements, as well as parameters related to the biological catalyst quality (yeast viability) and potential hazards due to microbial contamination (bacterial cell counts, fermented broth acidity). We termed this metric the Industrial Performance (**Methods**).

The correlation of strain clusters with different components of industrial performance (i.e. acidity and yeast viability) suggests that specific phenotypes are more detrimental to the industrial process. Organic acids are the main metabolites produced by *L. fermentum*, and is involved in the reduction of yeast viability and metabolic capacity due to intracellular acidification and anion accumulation^27^. Acidity is, therefore, an indirect measurement of its metabolism, and its negative impact on ethanol yield and industrial performance demonstrates that the metabolism and growth of bacteria is critical for this industry.

For both mills, a similar trend was observed in which poorer industrial performances are observed when strain clusters 1 and 3 are dominant in the *L. fermentum* population (*>*50% in relative abundance). In contrast, dominance of *L. fermentum* strain cluster 2 is linked with improved industrial performance (p < 0.02 in both mills; Wilcoxon rank-sum test, **Figure 3A**). This new evidence suggests that, contrary to what is currently considered in the literature, the impact of *L. fermentum* in the fermentations is driven by its population’s strain composition, rather than its abundance in the process^19,28–32^. It also suggests that current contamination control practices need to take into account the strain dominance, in order to choose the best molecule or mode of application. This can only be achieved with higher resolution diagnostics.

**Figure 3:**
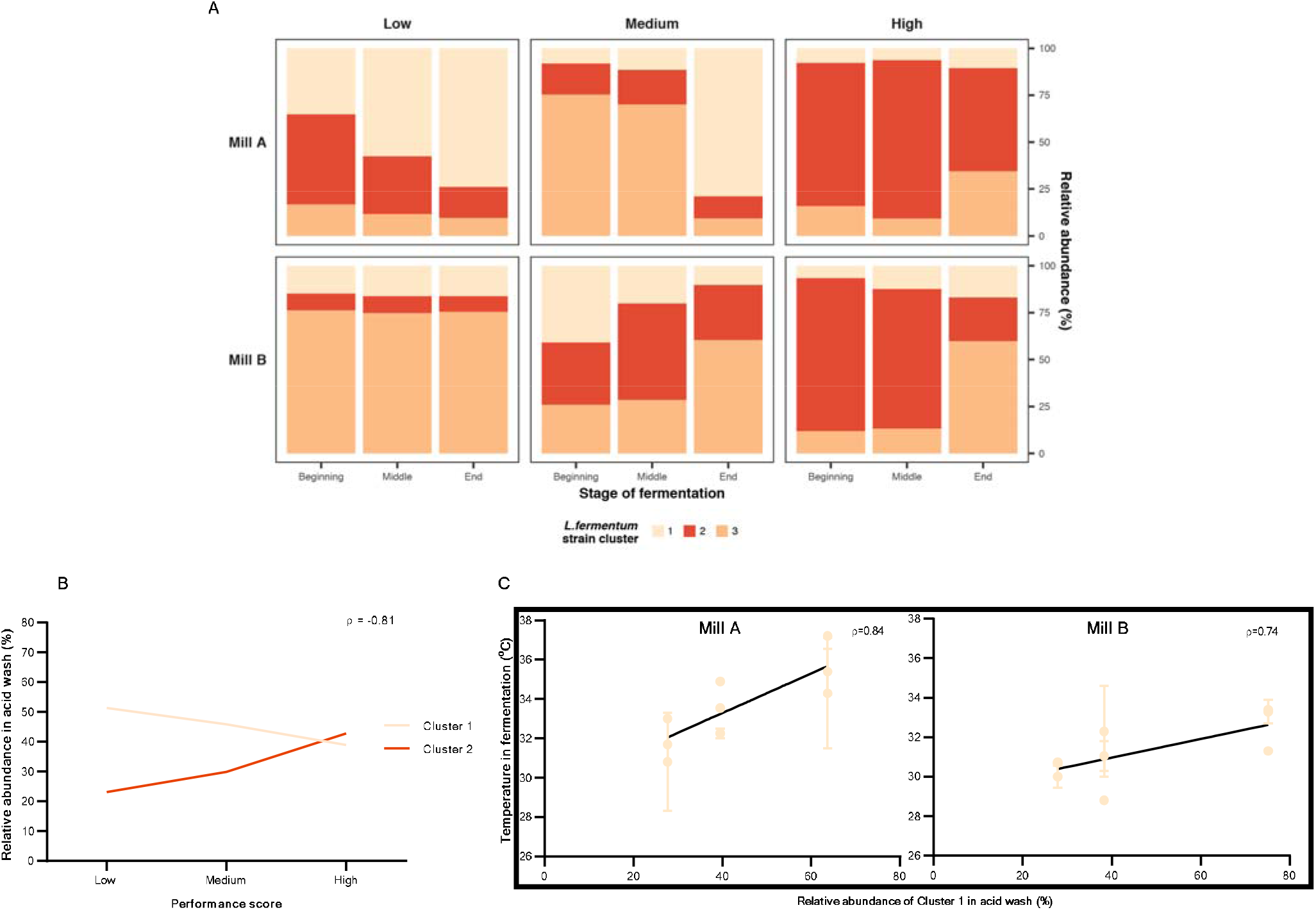
*L. fermentum* strain dominance is associated with process performance. **A:** The mean relative abundances of the 3 *L. fermentum* strain clusters, within its population, in all fermentation steps (beginning, middle and end) against industrial performance score (Low, Medium and High). Strain cluster 2 dominates high-performance batches during most of the fermentation process (p<0.02), whereas clusters 1 and 3 reach their lowest relative abundances in these high-performance batches. **B:** The interplay between *L. fermentum* strain clusters 1 and 2 in the acid wash are intimately connected the population composition in the beginning of the fermentation (ρ = 0.52 and FDR = 0.021, ρ = 0.82 and FDR = 0.004 for clusters 1 and 2, respectively) and with performance scores. The competition between these two strain clusters decides the fate of the next fermentation batch, being mutually exclusive in the acid wash (ρ = −0.81, FDR = 1.6 x 10^-4^). **C:** Higher fermentation temperatures privilege the detrimental strain cluster 1, being directly correlated with its higher relative abundance in the acid wash, and in subsequent fermentations, for both mills (ρ = 0.84, FDR = 0.0048 for mill A; ρ=0.74, FDR= 0.0185 for mill B).

To understand the mechanism underlying these different population composition in fermentation with high and low industrial performance we analysed the *L. fermentum* population composition for each unitary step of the fermentation process. The population composition during the beginning of the fermentation is crucial for defining the industrial performance of the fermentation batch. More specifically, the relative abundance of strain cluster 2 is directly correlated with higher industrial performances scores (ρ = 0.73, FDR = 0.003), and the abundance of strain clusters 1 and 3 are linked to lower industrial performance scores (ρ = −0.51, FDR = 0.042; ρ= −0.70, FDR=0.006, respectively) (**Supplementary Table 6**).

Notably, the strain level composition of strain clusters 1 and 2 in the acid wash tank, the unitary step immediately prior to the new fermentation batch, is directly correlated with their relative abundance in the beginning of the fermentation defining the strain composition of new batches (ρ = 0.60 and FDR = 0.016 for cluster 1; ρ = 0.85 and FDR = 0.0002 for cluster 2) (**Supplementary Table 6**). The direct correlation between acid wash and the beginning of the fermentation suggests that the microbial community composition is mainly driven by cell recycle, rather than the addition of novel contaminants through the broth. This hypothesis is further backed by the higher dissimilarity observed in the broth, when compared to the other fermentation steps (**Supplementary Table 1**), suggesting a more diverse and different community composition than the one found in actual fermentations.

The population dynamics of the two strain clusters suggest direct competition in the acid wash (ρ = −0.81, FDR = 1.6 x 10^-4^; **Figure 3B**). The outcome of this competition between closely related strain clusters in the acid wash is decisive for the industrial performance of the following fermentations.

Focusing on identifying potential process parameters that could influence this strain level dynamics we have analysed the correlation between the strain level compositions of *L. fermentum* populations against all registered metadata. We have also broken down this analysis into the specific operational processes, in order to identify any trends correlated with a specific step of the ethanol production process. The fermentation temperature found in individual vessels is the key driving factor defining *L. fermentum* strain level composition in the acid wash. More specifically, higher temperatures throughout the fermentation lead to a higher relative abundance of strain cluster 1 in the acid wash, after the fermentation ρ = 0.55, FDR = 0.0001; **Figure 3C**), favouring this specific cluster. In that way, the different fermentation batches are intimately connected, and the impact on process is more dependent on the established microbiome rather than novel contaminants entering via fermentation broth.

Temperatures above 32°C during the fermentation process are correlated with higher relative abundances of cluster 1, and with lower performance scores. Keeping the fermentation below this temperature threshold would, therefore, favour strain cluster 2, a less detrimental cluster of *L. fermentum*.

To test if our hypothesis is correct we sought to investigate if there are strain level variance in the temperature dependence of the growth rate of different Lactobacillus strains. In laboratory conditions it was also possible to replicate this phenomenon. The growth rate (h^-1^) of actual industrial lactobacilli isolates was compared in two different temperatures (30°C and 37°C). While *L. amylovorus* shows a growth rate 14% higher at 30°C, the three different strains of *L. fermentum* are favoured by higher temperatures, but with considerable differences among them. Strain F1 had its growth rate leaping from 0.03 to 0.21, a 5 fold increase in the growth rate. Strain F2 shows a growth rate 76% higher at 37°C, and Strain F3 a growth rate 170% higher (**Figure 4**). These results suggest that the strain level dynamics of microbial populations is directly affected by environmental factors, and might be responsible for shaping the functionalities of microbiomes.

**Figure 4:**
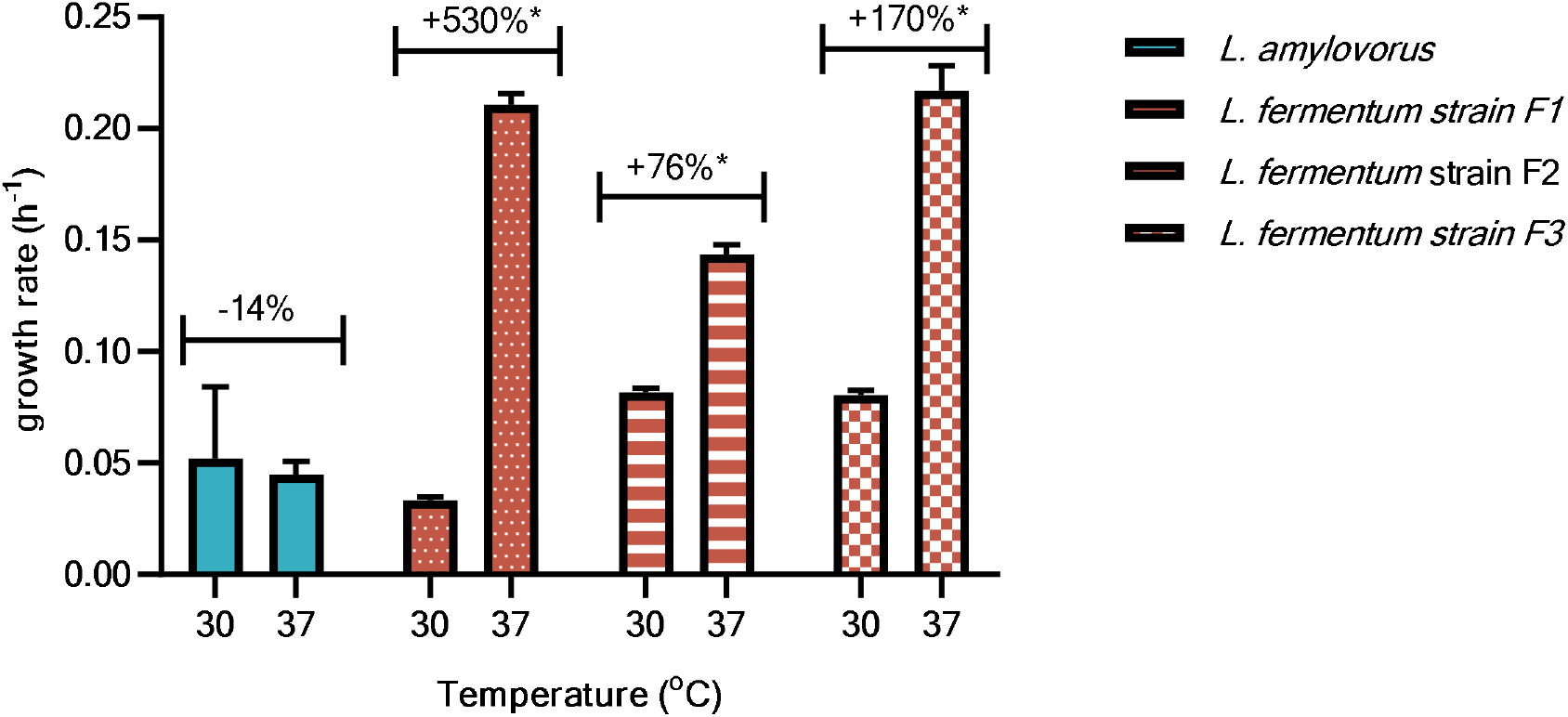
The influence of temperature in the growth rate (h^-1^) of different industrial isolates. Higher fermentation temperatures favours *L. fermentum* in detriment of *L. amylovorus*, which has its growth rate reduced in 14% (from 0.052±0.026 to 0.045±0.005). Within *L. fermentum*, different strains display diverse responses to the increase of temperature. Strain F1 shows an average increase in its growth rate of 530% (from 0.33±0.001 to 0.211±0.004). Strain F2 grows 76% faster (from 0.082±0.002 to 0.143±0.004), and strain F3, 170% (from 0.08±0.002 to 0.217±0.009). *p < 0.05.

The comparison of the pangenomes of the 3 *L. fermentum* strain clusters revealed the presence of functions and pathways not found in the other strains, which may be associated with their impact and prevalence on the industrial process. (**Methods, Supplementary Table 7**). Strain clusters 1 and 3 (the most detrimental ones) present several unique genes which seem to be correlated to an adaptation for living and growing in the fermentation environment. Furthermore, strain cluster 1 contains arginine biosynthesis genes (KO1438), an amino acid that *L. fermentum* is otherwise known to be auxotrophic for^34^. Ameliorating this auxotrophy could improve the competitive fitness of this strain in an amino acid depleted environment^35^.

### Loss of performance in fermentation is related to the competition between bacteria and yeast, and the metabolite profile of *L. fermentum* strains

To elucidate the impact of strain variation on the ethanol fermentation yield by *S. cerevisiae*, we performed static batch cultivations, using the model yeast strain PE-2 and *L. fermentum* strains isolated from the samples used in our microbiome analyses (**Figure 5A**). Here, we conducted pairwise fermentations that simulated the conditions of a typical industrial setup by applying a yeast-to-bacteria ratio of 100:1^5^, and using a chemically-semi defined synthetic medium that resembles the sugarcane molasses-based broth^36,37^.

**Figure 5:**
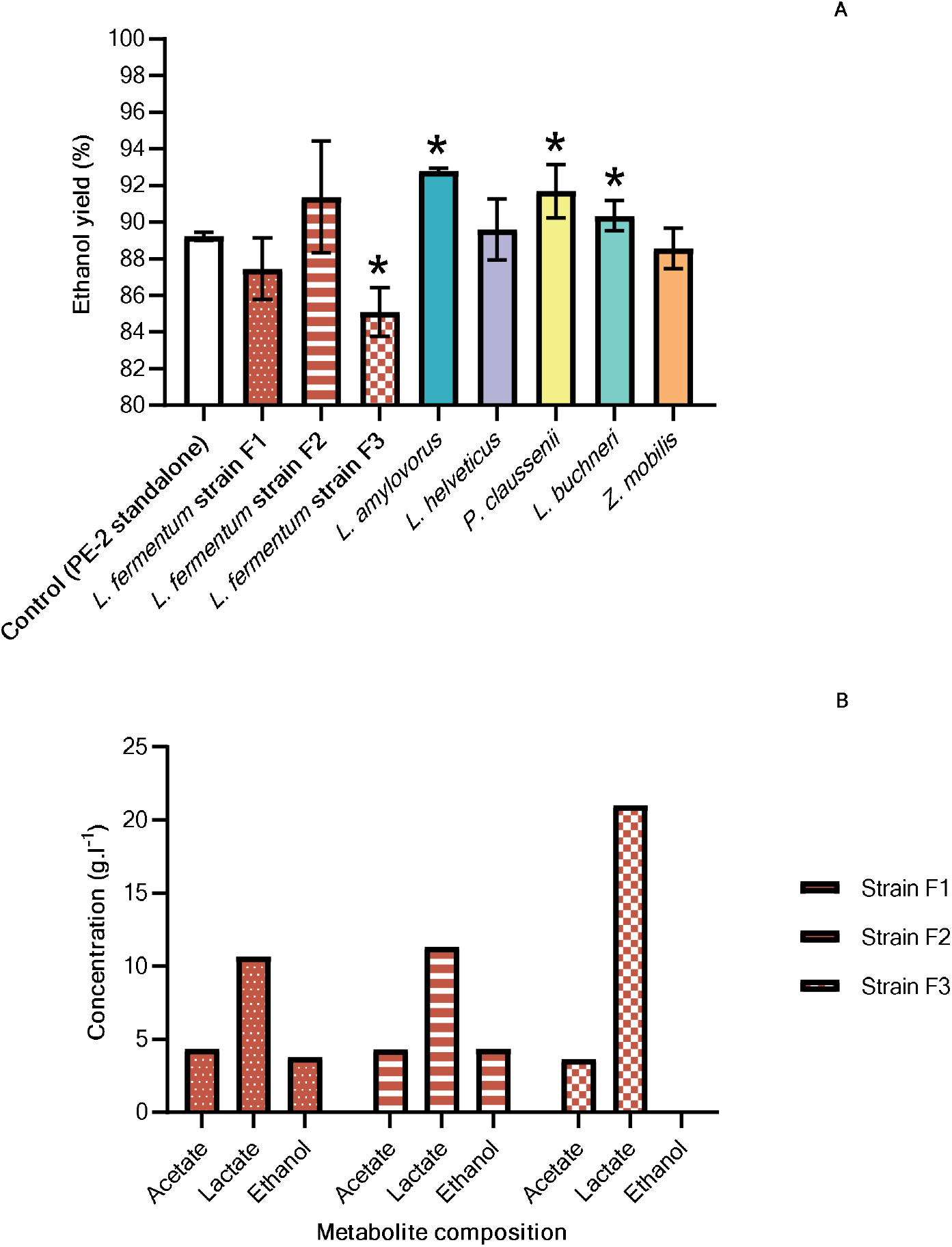
Specific *L. fermentum* strains cause reduction in bioethanol fermentation yield. **A:** Ethanol yield of control fermentations (PE-2 yeast strain standalone) is compared against pairwise fermentations with yeast strain PE-2 and 3 *L. fermentum* industrial isolates (filled bars); and with each of the 5 most common contaminant bacteria found in sugarcane ethanol fermentations. Of these species, which account for 80% of the contaminant microbial community, only *L. fermentum* strains F1 and F3 reduce ethanol yield, albeit only F3 being statistically significant (by 1.98±1.68% and 4.63±1.35% respectively; multiple t-test). *p < 0.05. **B:** The metabolite profile of the different *L. fermentum* strain isolates supernatants. Strains F1 and F2 show a similar metabolite production. On the other hand, strain F3 – the most detrimental one – presents a remarkably different metabolite production, with twice as much lactate (up to 21 g x l^-1^) and no ethanol produced, when compared to the other strains. This suggests that the mode of inhibition of lactobacilli is directly correlated with their organic acid production profile^6^.

The yield obtained from the standalone fermentation by yeast PE-2 served as a control. To further contextualise our findings, we repeated these experiments on the 6 most abundant bacterial species (**Figure 1B**). Altogether, the species we analysed account for almost 80% of known species in the contaminant microbiome (**Supplementary Table 4**).

Our results demonstrate that the presence of *L. fermentum*, compared to other abundant bacterial species, can have a negative impact on fermentation performance^20^, with one strain decreasing ethanol yield by 4.63±1.35% (**Figure 5A**). This contrasts with *L. amylovorus, P. claussenii* or *L. buchneri*, which showed positive effects on ethanol yield (multiple t-test, p < 0.05). Both *L. amylovorus* and *P. claussenii* have a homofermentative metabolism, which has been shown to be less detrimental to *S. cerevisae* in this fermentation setup^6^. Notably, *L. buchneri* strains have previously been shown to produce considerable amounts of ethanol and lactate from glucose^38^. This ethanol production might contribute to greater ethanol titres and yields, despite competing with yeast during the fermentation process. These observations are reinforced by our metagenomics analyses, where the presence of these species correlated poorly with acidity, which was strongly associated with lower ethanol yields (**Figure 2B**).

Interestingly, we provide experimental evidence that not all *L. fermentum* strains are detrimental, as in the case of strain F2 (91.38±3.04%), which was comparable to control (89.23±0.23%; multiple t-test, p < 0.05). When comparing the metabolite profile of supernatants from monocultures of the *L. fermentum* isolate strains, we observe that there is a striking difference between the somewhat neutral strain F1 and the beneficial strain F2 with the detrimental strain F3. Not only its organic acid production titre is twice as the one observed for strains F1 and F2, it also does not produce ethanol, which will have a direct impact on final ethanol titre and yield of co-cultures with *S. cerevisae* (**Figure 5B**). These findings are in accordance to literature observations, which suggest that the ratio between different organic acids is more important than the high titres of specific organic acids for the inhibition of *S. cerevisae*^27,39^. This data demonstrates the importance to monitor not just the total acidity, but also the composition of the organic acid pool, in order to adapt the response to microbial contamination accordingly to this particular data.

In the detrimental strain F3 genome we identified a unique gene cluster that is involved in glycerol catabolism (KO2440, KO6120, KO6121 and KO6122). Glycerol is the second most abundant metabolite produced by *S. cerevisiae* in alcoholic fermentations^33^. The ability to use glycerol as an electron acceptor, as it is done by *L. reuteri*^40^, could likely provide a competitive advantage for these strains when growing in the presence of yeast, allowing them to exploit this exclusive niche created by yeast metabolism. Moreover, more of the carbon source could be deviated towards biomass production instead of energetic metabolism, since this strain would not require to reduce acetaldehyde into ethanol to rebalance the NADH/NAD^+^ pool, as it is commonly done in lactobacilli. The metabolite profile data indeed suggests that such strain lacks ethanol production under fermentative conditions and provides a mechanism for its particular detrimental effects on overall ethanol yields. In order to identify functions or pathways in our *L. fermentum* strains that may have contributed to their distinct fermentation behaviour, we sequenced their genomes and conducted a comparative analysis (**Methods, Supplementary Table 7**).The genetic diversity found between these strain clusters could allow, in theory, for the development of modern diagnostic tools (e.g. qPCR analysis) which could predict the performance of future fermentation batches, improving the process control of this industry.

## Discussion

Taken together, we have demonstrated that strain-level variation affects the output of a well-controlled industrial bioethanol fermentation process. This highlights the importance of using high-resolution, cross-sectional analysis of the contaminant microbiome in combination with relevant industrial indicators. We observe the interplay between *L. fermentum* and *L. amylovorus* that suggests competition between the two species in 2 ethanol producing mills. We also demonstrate the utility of the industrial sugarcane ethanol fermentation as a model system to study the dynamics and ecological interactions of microbial community, due to its highly compartmentalized setup, which enables the impact of perturbations (such as strain-level alterations) to be easily quantified.

Along these lines, we also introduce a novel metric by which to assess and compare the performance of an industrial bioethanol fermentation batch. Using this metric, we associated genetic variants of *L. fermentum* with fermentation efficiency, and identified the mechanism underlying the prevalence of detrimental strains as being the temperature of fermentations. Higher fermentation temperatures, above 32°C, privilege specific *L. fermentum* strains, which become the dominant strains in the population, and are considerably more detrimental to yeast fermentation, due to their different metabolic profile. Selective removal of the identified strain clusters could potentially improve ethanol yield by almost 5%, translating to estimated economic gains of 690 million USD^41^ and more than 2 million tons per year of CO_2_ emissions for the Brazilian bioethanol industry^42^.

## Acknowledgements

The authors would like to acknowledge Marcos Vinícius, Thais Granço and Rafael Alves on their support on acquiring industrial samples and metadata. The authors would like to acknowledge Prof Dr Adriano Azzoni and Prof Dr René Schneider on providing access to laboratory equipment for processing industrial samples. The authors would like to acknowledge the support of Georges Neto on the processing of samples from the first sampling timepoint batch.

The research was supported by funding from The Novo Nordisk Foundation under NFF grant number: NNF10CC1016517. SSL acknowledges support from EMBO (ALTF 137-2018). TOB acknowledges funding from FAPESP, with grant numbers: 2015/50684-9 and 2018/17172-2.

## Author contributions

FSOL, MAM, KK, SSL, TOB, GP and MOAS designed the study. FSOL, TOB and BLVC collected and processed industrial samples and metadata. FSOL prepared the metagenomics samples for sequencing. MAM, KK and SSL performed bioinformatics analyses. MAM, KK, SSL, FSOL, GP and MOAS critically analysed bioinformatics results. FSOL isolated industrial strains. FSOL designed and performed laboratory scale fermentations. FSOL and MOAS critically analysed the laboratory scale fermentation results. FSOL, MAM, KK, SSL and MOAS wrote the manuscript. All authors have provided inputs for the manuscript.

## Methods

### Chemicals

Unless stated otherwise, all chemicals and reagents used were purchased from Sigma-Aldrich (St. Louis, MO, USA).

### Sampling strategy

We sampled two independent ethanol mills (named Mill A and Mill B) in the production season of 2017. Both mills are located in the State of São Paulo, Brazil - in a region with the prevalence of the humid subtropical climate (*Cfa*) with an annual precipitation of around 2000 mm, and with a sea-level altitude of *ca*. 600m. The mills were completely independent from each other, with a distance greater than 300 km apart, and have raw material sourced from different producers and sugarcane fields. Both mills operated via fed-batch fermentations (Melle-Boinot setup), and had a similar ethanol production capacity with a daily output of ca. 400m^3^ of ethanol. Mill A was sampled in the dates: 26/05/2017; 26/10/2017 and 17/11/2017. Mill B was sampled in the dates: 02/06/2017; 29/10/2017 and 03/11/2017. The following steps of the ethanol production process were sampled: (1) Fermentation broth (Feeding line with fresh fermentation media); (2) start; (3) middle; (4) end of fermentation; (5) yeast cream after separation of the wine, which is sent to distillation centrifugation); and (6) biomass after acid wash treatment (sulphuric acid pH 2.5 for 1hour). The phases of the fermentation were defined according to the feeding regimen of each mill: the beginning was set as the beginning to the middle of the feeding; the middle was defined from the middle to the end of the feeding, and the end was defined as the final hours of fermentation, after the feeding had ceased. Due to the fact that different vessels are fed sequentially, we were able to sample different vats at different stages of fermentation, during the same sampling. Samples were collected directly from the production process and diluted 1x in a sterile Phosphate Buffered Saline (PBS) solution with glycerol (50%). The samples were readily frozen in dry ice, until final storage in ultrafreezer (−80°C). Each mill had several vessels operating in the same fermentation step, which allowed for process replicates. Samples were taken in duplicates.

### Industrial metadata

The industrial metadata was provided by the operational staff from each mill, and consisted on key process control parameters collected and registered by industrial staff, related to the ethanol fermentation. Those parameters were: Ethanol yield daily average); ethanol yield (weekly average); acidity from wine (g_acetic_ acid equivalent.l^-1^, where g_acetic_ acid equivalent is related to the amount, in g.l^-1^, of acetic acid equivalent obtained via titration); yeast cell counts in the fermentation (CFU); bacteria cell counts in the fermentation (CFU); yeast viability (% of the population); yeast budding rate (% of the population); vessel current volume (in m^3^); vessel operational status (idle, feeding, running or finished fermentation) and vessel temperature (in °C).

For correlation analyses, the data was converted into monthly averages.

### DNA extraction and sequencing of isolates and metagenomes

All DNA extractions were performed using the DNeasy Powerlyzer Powersoil Kit (QIAGEN, Hilden, Germany), according to manufacturer’s instructions. Pure lactobacilli isolates had their DNA extracted using MasterPure™ Gram Positive Purification Kit (LGC Biosearch Technologies, Hoddesdon, UK). All DNA extraction quantifications were performed with Qubit Fluorometer (Thermo Fischer Scientific, Waltham, MS, USA). Due to low biomass content, DNA extraction was not possible for broth sample (Process Step 1) from Mill A at the 3^rd^ sampling timepoint. Shotgun metagenomics and isolates genome sequencing was performed on the NextSeq 500 using NextSeq High Output v2 Kit (300 Cycles) (Illumina, San Diego, CA, USA) by the Sequencing Core Facility at The Novo Nordisk Foundation Center for Biosustainability (Technical University of Denmark, Kongens Lyngby, Denmark). The library preparation was performed using the KAPA HyperPlus Library Prep Kit (Roche, Basel, Switzerland), and the indexing kit used was the Dual Indexed PentAdapters, Illumina compatible (PentaBase, Odense, Denmark). Quantity and quality control were performed using Qubit dsDNA HS Assay Kit (Invitrogen, Carlsbad, CA, USA) and DNF-473 Standard Sensitivity NGS Fragment Analysis Kit (1 bp −6000 bp; Agilent, Santa Clara, CA, USA). Average library length was 341 bp. The sequencing reads length were 150 base pair paired-end (2×150 bp). The index (i7 and i5) reads were 8 bp, dual indexed and flow cell loading was 1.3 pM. The sequencing chemistry used was 2-channel sequencing-by-synthesis (SBS) technology, and Phix control V3 (Illumina San Diego, CA, USA) was added (2.5%).

### Processing of genomic and metagenomic data

Raw reads (from both metagenomic and isolate sequencing) underwent quality trimming, i.e. filter out adapter and universal primer sequences, as well as low quality bases (< Q20), reads shorter than 75 bp and duplicated reads (**Supplementary Table 2**), as previously described^43^.

All 481 *S. cerevisiae* genomes from NCBI genome database (August 2018) were downloaded. Reads were aligned to concatenated genomes using the BWA mem model with default parameters^43^. Reads over 95% identity were considered to belong to *S. cerevisiae* (SC reads) and not used in our analyses (**Supplementary Table 2)**. Kraken^44^ was selected for taxonomic read assignment of non-SC reads as it has been shown to perform well in benchmarking studies^45,46^, especially for medium-low complexity microbiome communties^47^ such as the ones in industrial fermenters. Specifically, Kraken v. 0.10.5-beta was applied on non-yeast reads with default settings against the minikraken 2017.10.18 8GB database. Bracken^48^ v. 2.0 was used for accurate species abundance estimation with parameters-r (read length) 150 and-l (level) S (species).

### Analyses of metagenomics data

Rarefaction of read counts and subsequent analyses were done using R package *vegan*^49^. We considered the 10 most prevalent contaminant species as those with the highest median relative abundance across all samples. Microbial community compositions were compared using Bray-Curtis distance on species relative abundance and Permutational Multivariate Analysis of Variance (PERMANOVA) with 999 permutations and the Bray-Curtis method was applied by providing Mill/Process step/Date as function. Pairwise Spearman’s correlation coefficient was calculated for pairs of metadata variables, and between metadata variables and taxon abundances. False discovery rate (FDR) was calculated using Benjamini-Hochberg (BH) method, with FDR < 0.05 used as cut-off.

### Strain level profiling

To profile the strains within a bacterial species, we used a published pipeline^43,50^ with the following modifications: (1) all contigs from one NCBI genome were concatenated as a consecutive sequence with spacers (N of 100bp); (2) genome assemblies from the NCBI database were automatically downloaded and renamed; (3) the new NCBI accession ID system in place of the old sequence ID and taxonomic ID system, which allowed more assemblies to be included. For each species, genomes were downloaded from the NCBI Genome database (August 2018) to construct a SNV and pan-genome database. In cases where more than 200 genome assemblies existed, only complete assemblies and chromosome-level assemblies were used. The SNV-based core-genome strain profiling was used for downstream analyses. Strains with a maximum relative abundance less than 5% in at least one sample, or present in less than 20% samples were discarded.

For each species, Spearman’s correlation coefficient was calculated among different strains’ relative abundance of all metagenomic samples. False discovery rate (FDR) was calculated using Benjamini-Hochberg (BH) method. FDR < 0.05 was used as the significant level cut-off. Strain abundance dissimilarity among different samples was calculated by Euclidean distances and hierarchical clustering was performed. As a result, all strains were clustered into two to six strain clusters for each species, where no significant negative correlation could be captured in each strain cluster (see **Supplementary Figure S3** as an example from *L. fermentum*).

### Industrial performance calculation

The industrial performance calculation was obtained by the product of the multiplication of the parameters directed correlated with process performance (i.e. ethanol yield and yeast viability), divided by the product of the multiplication of the parameters inversely correlated with process performance (i.e. bacterial cell counts and acidity titre):

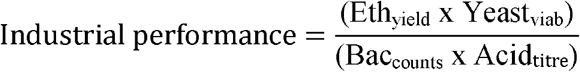

#### Equation 1: Proposed equation for obtaining a general industrial performance score

The score is obtained by multiplying ethanol yield (Eth_yield_) and yeast viability (Yeast_viab_) values, and dividing its product by the product obtained from the multiplication of bacterial cell counts (Bac_counts_) and wine acidity titre (Acid_titre_) values.

### Strains used in laboratory experiments

*Saccharomyces cerevisiae* strain PE-2 was kindly provided by Prof. Thiago Olitta Basso. Strains of *Lactobacillus amylovorus* and *Lactobacillus fermentum* were isolated from stored industrial samples. Strains of *Pediococcus claussenii, Lactobacillus helveticus, Lactobacillus buchneri* and *Zymomonas mobilis* were purchased from ATCC (Manassas, VA, USA).

### Isolation and maintenance of industrial strains

For strain isolation, a previously introduced protocol was used^13^. Briefly, industrial samples were serially diluted in sterile PBS and plated in Man Rogosa Sharpe (MRS) Agar media, containing cycloheximide (0.1% v.v^-1^) in order to inhibit yeast growth. Plates were incubated at either 30°C or 37°C statically. A loopful of an isolated colony was grown in liquid MRS in the same conditions, and stored at −80°C (see section “DNA extraction and analysis of bacterial isolates”). Yeast strains were cultured in Yeast Potato Dextrose (YPD) media, at 30°C. Lactobacilli were cultured in MRS media, either at 30°C or 37°C, and *Zymomonas mobilis* was cultured at Trypsin Soy Broth (TSB) media, at 30°C. All cultivations were performed statically, in *ca*. 5ml volume.

### DNA extraction and analysis of bacterial isolates

Pure isolates were grown overnight in adequate media and conditions, as mentioned in the previous section. After growth, cells were pelleted via centrifugation (> 10,000g for 4 min.) and their genomic DNA was extracted using the MasterPure^™^ Gram Positive DNA Purification Kit (Lucigen Corporation, Middleton, WI), according to manual’s instruction.

### Bacterial isolate assembly

A *de Bruijn* graph-based assembler, SPAdes 3.12^51^, was used for the genome assembly of bacterial isolates, using the following parameters: “-m 300-k 33,55,77,99,127”. To complement the *de novo* assembly, reference-assisted genome assembly was performed with idba_hybrid (v 1.1.1)^52^ with the following parameters “--pre_correction --mink 120 --maxk 180 --step 10 -- min_contig 300 --reference [the reference genome downloaded from NCBI for each species]”. Two modifications were made in the source code before compiling IDBA_UD: in file src/basic/kmer.h constant kNumUint64 was changed from 4 to 8 to allow maximum kmer length beyond 124; in file src/sequence/short_sequence.h constant kMaxShortSequence was set to 512 to support longer read length. Final assembly results were summarized in **Supplementary Table 4, Methods**.

### Genome-based functional analysis

ORFs were predicted on assembled strain genomes using MetaGeneMark v.3.26^53^. Predicted proteins were annotated using eggNOG mapper^54^ with the following settings: mapping mode: DIAMOND, automatic taxonomic scope, orthologs: restrict to one-to-one (prioritize precision), GO evidence: use experimental-only terms (prioritize quality). From the resulting file, KEGG ortholog ids were extracted by significant matches (e-value < 10^-5^) and those that were unique for each strain cluster were characterized in terms of KEGG pathway/module membership (KEGG Mapper Pathway Reconstruct)^55^.

### Fermentation experiments

Fermentations were performed in 96 deep-well plates, with either pairwise cultivations (yeast:bacteria at a 100:1 ratio)^8^, or standalone yeast or bacteria cultivations. The media used is a semi-synthetic media, able to simulate sugarcane molasses based media (SM)^37^. Briefly, all strains were cultured in their optimal media and conditions (see “Strains” and “Isolation of industrial strains and maintenance” sections), for up to 48h. After that, the biomass was calculated via optical density (OD; 600 nm wavelength). All cells were pelleted via centrifugation (3400 x g, 4°C, 15 min) and washed twice with sterile PBS. Subsequently, cells were diluted in SM diluted in sterile Milli-Q H2O (10x, final sugar concentration of 18g.l^-1^) for an OD value of 1.0. Strains were later diluted in fresh SM media in specific wells in the 96 deep-well plate to a final OD value of 0.1.

The lactobacilli growth rate analysis was performed at 30 and 37°C, under agitation (double orbital, fast mode) in Synergy H1 plate readers (Biotek Instruments, Inc. Winooski, VT, USA). OD was measured every 30 minutes for 24h. The growth rate was later calculated using the R package *growthcurver*^56^.

All the pairwise cultivations were performed statically, overnight, at 30°C, in ca. 1 ml volume. The fermentations were performed in triplicate. The carbohydrate titre and composition (sucrose, glucose and fructose) and fermentation metabolites (glycerol, ethanol, and acetic acid) were determined by high-performance liquid chromatography (HPLC) (UltiMate 3000, Thermo-Fischer Scientific, Waltham, Massachusetts, USA). The analites were separated using an Aminex HPX-87H ion exclusion column (Bio-Rad, Hercules, California, USA) and were isocratically eluted at 30°C, with a flow rate of 0.6 ml.min^-1^, using a 5mM sulphuric acid solution as mobile phase. The detection was performed refractrometrically.

Ethanol yield was calculated according using the following equation:

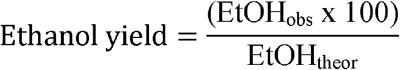

#### Equation 2: Ethanol yield calculation

Where: EtOH_obs_ = the observed ethanol titre on each sample. EtOH_theor_ = the maximum theoretical ethanol titre for each sample. Obtained by multiplying the sugar titre from the broth solution with the stoichiometric conversion factor for ethanol production (i.e. 0.5111) ^57^.

Community composition was resolved via flow-cytometry (BD LSRFortessa^™^, BD Biosciences, Franklin Lakes, New Jersey, USA). A sample from each well (10 μl) was taken after the overnight cultivation, and was transferred to a new microplate and diluted in 190μl PBS buffer (pH 7.4). Yeast and bacteria populations were resolved via front and side scatter comparison (SSC versus FSC). The statistical analyses were performed using the software GraphPad Prism 8. The difference on final ethanol yield was analysed by multiple t-tests (statistical significance analysis with alpha value of 0.05).

Lactobacilli supernatant metabolite profile was analysed via HPLC after 48h of growth, using the aforementioned analytical method. A pre-inoculum of lactobacilli stored at −80°C was grown in MRS for 24h. After that the OD from these cultures was measured and fresh MRS media was inoculated with a fixed OD of 0.1 and incubate statically at 37°C. After growth, the cells were separated via centrifugation and the supernatant was sent for further analysis.

### Metagenomic co-assembly and functional annotation

To overcome the imbalance between the sequencing yields of bacterial and fungal reads in different samples, achieve higher completeness of pan-genome regions with low sequencing coverage, and perform sample-wise gene presence and absence comparisons, co-assembly was performed for the non-SC reads. Non-SC reads were concatenated separately from all sequenced samples and the maximum *k*-mer depth was normalized to 100 fold by BBnorm (https://sourceforge.net/projects/bbmap/) before co-assembly. IDBA_ud (v. 1.1.1)^52^ was used for the assembly using the following parameters: “--min_contig 300 --mink 50 --maxk 124 --step 10 --pre_correction”. Co-assembly results are summarized in **Supplementary Table 3, Methods**. For the non-SC assembly, MetaGeneMark v. 3.26 was adopted to predict the coding DNA sequence (CDS) regions in the assembled metagenome contigs using the default parameters.

## References

1. Silvério da Silva, S. & Chandel, A. K. Biofuels in Brazil. (Springer, 2014). doi:10.1007/978-3-319-05020-1

2. Wang, M., Han, J., Dunn, J. B., Cai, H. & Elgowainy, A. Well-to-wheels energy use and greenhouse gas emissions of ethanol from corn, sugarcane and cellulosic biomass for US use. Environ. Res. Lett. 7, 13 (2012).

3. Wheals, A. E., Basso, L. C., Alves, D. M. G. & Amorim, H. V. Fuel ethanol after 25 years. Trends Biotechnol. 17, 482–487 (1999).

4. Basso, L. C., Basso, T. O. & Rocha, S. N. Ethanol Production in Brazil▢: The Industrial Process and Its Impact on Yeast Fermentation. Biofuel Prod. Recent Dev. Prospect. 1530, 85–100 (2011).

5. Amorim, H. V. et al. Scientific challenges of bioethanol production in Brazil. Appl. Microbiol. Biotechnol. 91, 1267–1275 (2011).

6. Basso, T. O. et al. Homo-and heterofermentative lactobacilli differently affect sugarcane-based fuel ethanol fermentation. Antonie van Leewenhoek 105, 169–177 (2014).

7. Lopes, M. L. et al. Ethanol production in Brazil: a bridge between science and industry. Brazilian J. Microbiol. 47, 64–76 (2016).

8. Ceccato-Antonini, S. R. Conventional and nonconventional strategies for controlling bacterial contamination in fuel ethanol fermentations. World J. Microbiol. Biotechnol. 34, 80 (2018).

9. Stubbendieck, R. M., Vargas-Bautista, C. & Straight, P. D. Bacterial Communities: Interactions to Scale. Front. Microbiol. 7, 1234 (2016).

10. Sinsabaugh, R. L., Manzoni, S., Moorhead, D. L. & Richter, A. Carbon use efficiency of microbial communities: stoichiometry, methodology and modelling. Ecol. Lett. 16, 930–939 (2013).

11. Ait Mueller, J. et al. Interspecies interactions are an integral determinant of microbial community dynamics. Front. Microbiol. 6, 1–11 (2015).

12. Makanjuola, D. B., Tymon, A. & Springham, D. G. Some effects of lactic acid bacteria on laboratory-scale yeast fermentations. Enzyme Microb. Technol. 14, 350–357 (1992).

13. Lucena, B. T. L. et al. Diversity of lactic acid bacteria of the bioethanol process. BMC Microbiol. 10, 298 (2010).

14. Bischoff, K. M., Liu, S., Leathers, T. D., Worthington, R. E. & Rich, J. O. Modeling bacterial contamination of fuel ethanol fermentation. Biotechnol. Bioeng. 103, 117–122 (2009).

15. Skinner, K. A. & Leathers, T. D. Bacterial contaminants of fuel ethanol production. J. Ind. Microbiol. Biotechnol. 31, 401–408 (2004).

16. Beckner, M., Ivey, M. L. & Phister, T. G. Microbial contamination of fuel ethanol fermentations. Lett. Appl. Microbiol. 53, 387–394 (2011).

17. Bonatelli, M. L., Quecine, M. C., Silva, M. S. & Labate, C. A. Characterization of the contaminant bacterial communities in sugarcane first-generation industrial ethanol production. FEMS Microbiol. Lett. 364, 1–8 (2017).

18. Costa, O. Y. A. et al. Microbial diversity in sugarcane ethanol production in a Brazilian distillery using a culture-independent method. J. Ind. Microbiol. Biotechnol. 42, 73–84 (2015).

19. Narendranath, N. V., Hynes, S. H., Thomas, K. C. & Ingledew, W. M. Effects of lactobacilli on yeast-catalyzed ethanol fermentations. Appl. Environ. Microbiol. 63, 4158–4163 (1997).

20. Rich, J. O., Leathers, T. D., Bischoff, Kenneth, M., Anderson, A. M. & Nunnally, M. S. Biofilm formation and ethanol inhibition by bacterial contaminants of biofuel fermentation. Bioresour. Technol. 196, 347–354 (2015).

21. Yang, C. et al. Strain-level differences in gut microbiome composition determine fecal IgA levels and are modifiable by gut microbiota manipulation. bioRxiv 544015 (2019). doi:10.1101/544015

22. Koch, C., Müller, S., Harms, H. & Harnisch, F. Microbiomes in bioenergy production: From analysis to management. Curr. Opin. Biotechnol. 27, 65–72 (2014).

23. Leathers, T. D. et al. Inhibitors of biofilm formation by biofuel fermentation contaminants. Bioresour. Technol. 169, 45–51 (2014).

24. Li, Q., Heist, E. P. & Moe, L. A. Bacterial Community Structure and Dynamics During Corn-Based Bioethanol Fermentation. Microb. Ecol. 71, 409–421 (2016).

25. Andrietta, S. R. R., Andrietta, M. G. S. & Bicudo, M. H. P. Comparação do rendimento fermentativo utilizando diferentes metodologias de cálculo para avaliação do desempenho de um processo industrial. STAB 30, 41–49 (2012).

26. Bhalla, A., Bischoff, K. M. & Sani, R. K. Highly thermostable xylanase production from a thermophilic Geobacillus sp. strain WSUCF1 utilizing lignocellulosic biomass. Front. Bioeng. Biotechnol. 3, 1–8 (2015).

27. Piper, P., Calderon, C. O., Hatzixanthis, K. & Mollapour, M. Weak acid adaptation: the stress response that confers yeasts with resistance to organic acid food preservatives. Microbiology 147, 2635–2642 (2001).

28. Liu, M. et al. Bacteriophage application restores ethanol fermentation characteristics disrupted by Lactobacillus fermentum. Biotechnol. Biofuels 8, 132 (2015).

29. Bassi, A. P. G., Meneguello, L., Paraluppi, A. L., Sanches, B. C. P. & Ceccato-Antonini, S. R. Interaction of Saccharomyces cerevisiae-Lactobacillus fermentum-Dekkera bruxellensis and feedstock on fuel ethanol fermentation. Antonie Van Leeuwenhoek 111, 1661–1672 (2018).

30. Reis, V. R. et al. Effects of feedstock and co-culture of Lactobacillus fermentum and wild Saccharomyces cerevisiae strain during fuel ethanol fermentation by the industrial yeast strain PE-2. AMB Express 8, 1–11 (2018).

31. Chang, I., Kim, B.-H., Shin, P.-K. & Lee, W.-K. Bacterial contamination and its effects on ethanol fermentation. J. Microbiol. Biotechnol. 5, 309–314 (1995).

32. Carvalho-Netto, O. V et al. Saccharomyces cerevisiae transcriptional reprograming due to bacterial contamination during industrial scale bioethanol production. Microb. Cell Fact. 14, 13 (2015).

33. Eli Della-Bianca, B., Olitta Basso, T., Ugarte Stambuk, B., Carlos Basso, L. & Karoly Gombert, A. What do we know about the yeast strains from the Brazilian fuel ethanol industry? Appl. Microbiol. Biotechnol. 97, 979–991 (2013).

34. Kuratsu, M., Hamano, Y. & Dairi, T. Analysis of the Lactobacillus metabolic pathway. Appl. Environ. Microbiol. 76, 7299–301 (2010).

35. Walford, S. Composition of cane juice. Proc. South African Sugar Technol. Assoc. 70, 265–266 (1996).

36. Basso, L. C., De Amorim, H. V., De Oliveira, A. J. & Lopes, M. L. Yeast selection for fuel ethanol production in Brazil. FEMS Yeast Res. 8, 1155–1163 (2008).

37. Lino, F. S. de O., Basso, T. O. & Sommer, M. O. A. A synthetic medium to simulate sugarcane molasses. Biotechnol. Biofuels 11, 221 (2018).

38. Liu, S., Skinner-Nemec, K. A. & Leathers, T. D. Lactobacillus buchneri strain NRRL B-30929 converts a concentrated mixture of xylose and glucose into ethanol and other products. J. Ind. Microbiol. Biotechnol. 35, 75–81 (2008).

39. Nicolaou, S. A., Gaida, S. M. & Papoutsakis, E. T. A comparative view of metabolite and substrate stress and tolerance in microbial bioprocessing: From biofuels and chemicals, to biocatalysis and bioremediation. Metab. Eng. 12, 307–331 (2010).

40. Stolz, P., Vogel, R. F. & Hammes, W. P. Utilization of electron acceptors by lactobacilli isolated from sourdough. Z. Lebensm. Unters. Forsch. 201, 402–410 (1995).

41. Portal Unica. Available at: http://www.unica.com.br/. (Accessed: 6th February 2019)

42. Börjesson, P. Good or bad bioethanol from a greenhouse gas perspective – What determines this? Appl. Energy 86, 589–594 (2009).

43. Kang, K. et al. The Environmental Exposures and Inner- and Intercity Traffic Flows of the Metro System May Contribute to the Skin Microbiome and Resistome. Cell Rep. 24, 1190–1202.e5 (2018).

44. Wood, D. E. & Salzberg, S. L. Kraken: ultrafast metagenomic sequence classification using exact alignments. Genome Biol. 15, R46 (2014).

45. Escobar-Zepeda, A. et al. Analysis of sequencing strategies and tools for taxonomic annotation: Defining standards for progressive metagenomics. Sci. Rep. 8, 12034 (2018).

46. Gardner, P. P. et al. Identifying accurate metagenome and amplicon software via a meta-analysis of sequence to taxonomy benchmarking studies. PeerJ 7, e6160 (2019).

47. Sczyrba, A. et al. Critical Assessment of Metagenome Interpretation—a benchmark of metagenomics software. Nat. Methods 14, 1063–1071 (2017).

48. Lu, J., Breitwieser, F. P., Thielen, P. & Salzberg, S. L. Bracken: estimating species abundance in metagenomics data. PeerJ Comput. Sci. 3, e104 (2017).

49. Dixon, P. VEGAN, a package of R functions for community ecology. J. Veg. Sci. 14, 927–930 (2003).

50. Oh, J. et al. Biogeography and individuality shape function in the human skin metagenome. Nature 514, 59–64 (2014).

51. Bankevich, A. et al. SPAdes: a new genome assembly algorithm and its applications to single-cell sequencing. J. Comput. Biol. 19, 455–77 (2012).

52. Peng, Y., Leung, H. C. M., Yiu, S. M. & Chin, F. Y. L. IDBA-UD: a de novo assembler for single-cell and metagenomic sequencing data with highly uneven depth. Bioinformatics 28, 1420–1428 (2012).

53. Zhu, W., Lomsadze, A. & Borodovsky, M. Ab initio gene identification in metagenomic sequences. Nucleic Acids Res. 38, e132 (2010).

54. Huerta-Cepas, J. et al. Fast Genome-Wide Functional Annotation through Orthology Assignment by eggNOG-Mapper. Mol. Biol. Evol. 34, 2115–2122 (2017).

55. Kanehisa, M., Furumichi, M., Tanabe, M., Sato, Y. & Morishima, K. KEGG: New perspectives on genomes, pathways, diseases and drugs. Nucleic Acids Res. 45, D353–D361 (2017).

56. Sprouffske, K. & Wagner, A. Growthcurver: An R package for obtaining interpretable metrics from microbial growth curves. BMC Bioinformatics 17, 1–4 (2016).

57. Raghavendran, V., Basso, T. P., da Silva, J. B., Basso, L. C. & Gombert, A. K. A simple scaled down system to mimic the industrial production of first generation fuel ethanol in Brazil. Antonie Van Leeuwenhoek 110, 971–983 (2017).

