## Supplementary Figures for "Strain dynamics of specific contaminant bacteria modulate the performance of ethanol biorefineries"

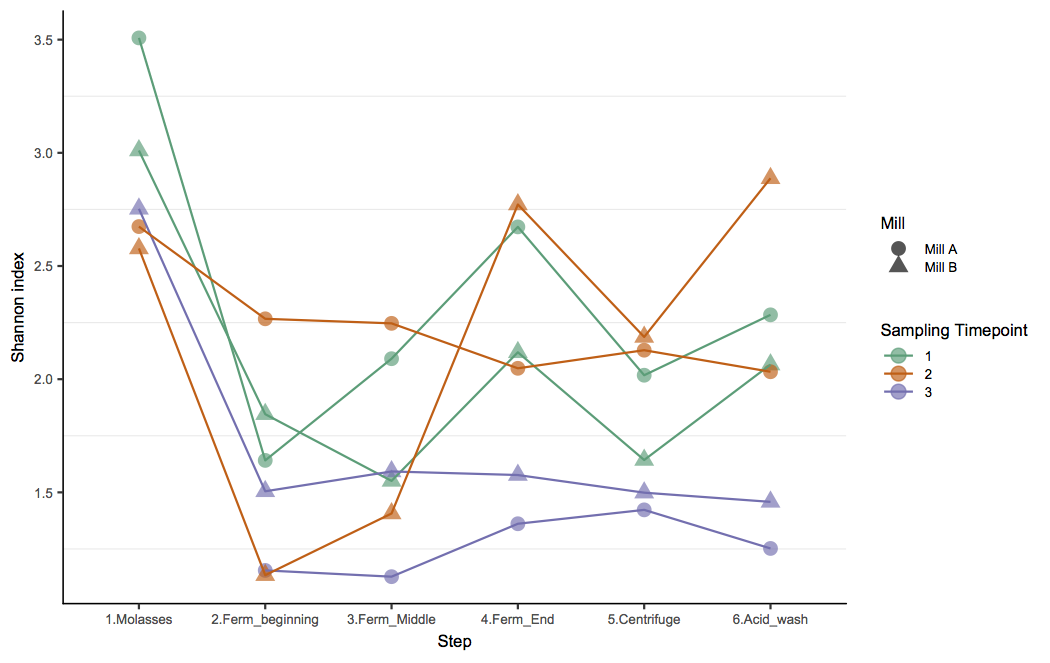

**Supplementary Figure 1. Alpha-diversity values from different process steps.** Average Shannon index values for samples from different process steps: 1) molasses (or broth); 2) Beginning of fermentation (Ferm_beginning); 3) Middle of fermentation (Ferm_Middle); End of fermentation (Ferm_End); 5) Centrifuge (or yeast cream) and 6) Acid wash. For all the 3 sampling timepoints the molasses (or broth) consistently shows higher Shannon index values, having a more uneven community composition structure.

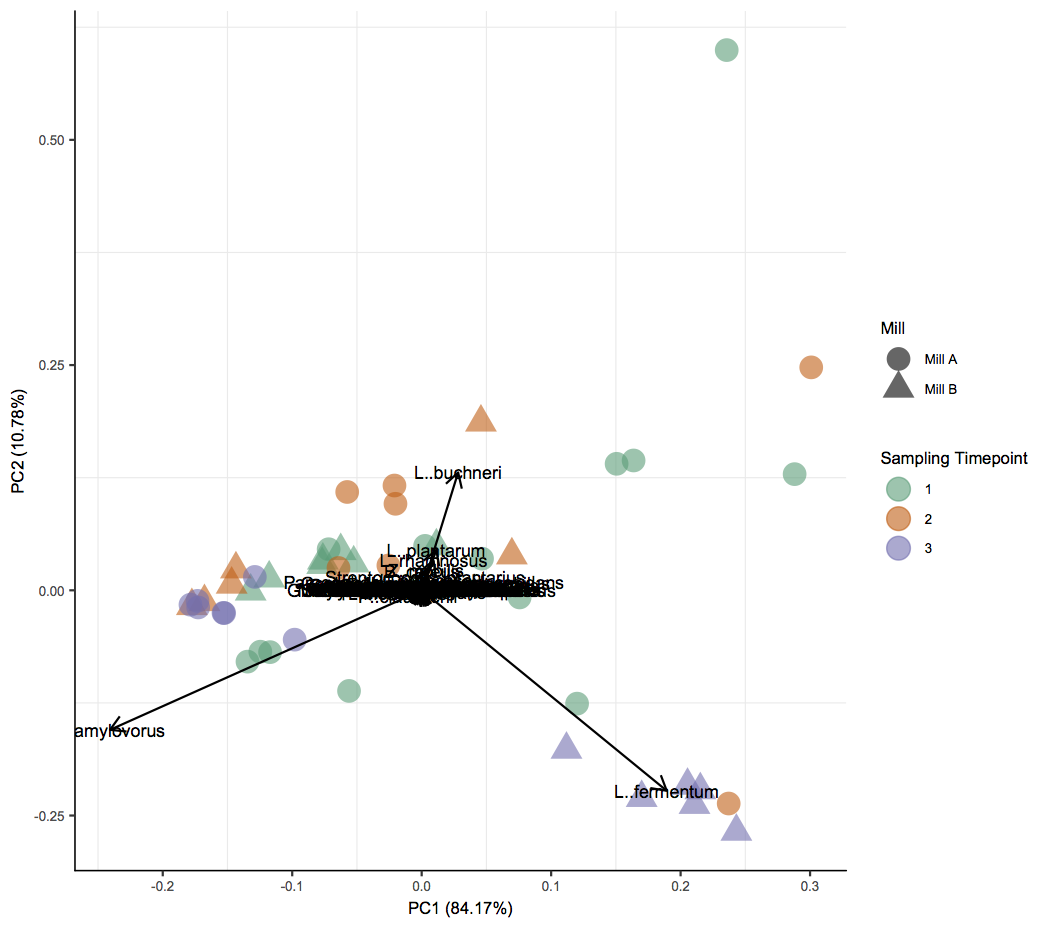

**Supplementary Figure 2**. **Principal component analysis of species-level relative abundances.** The main driving forces of community dissimilarity (PC1 = 84.08%) identified were the relative abundances of *L. fermentum* (right arrow), and *L. amylovorus* (left arrow). Shapes denote mills (circles for Mill A, triangles for Mill B) and colors denote sampling timepoint.

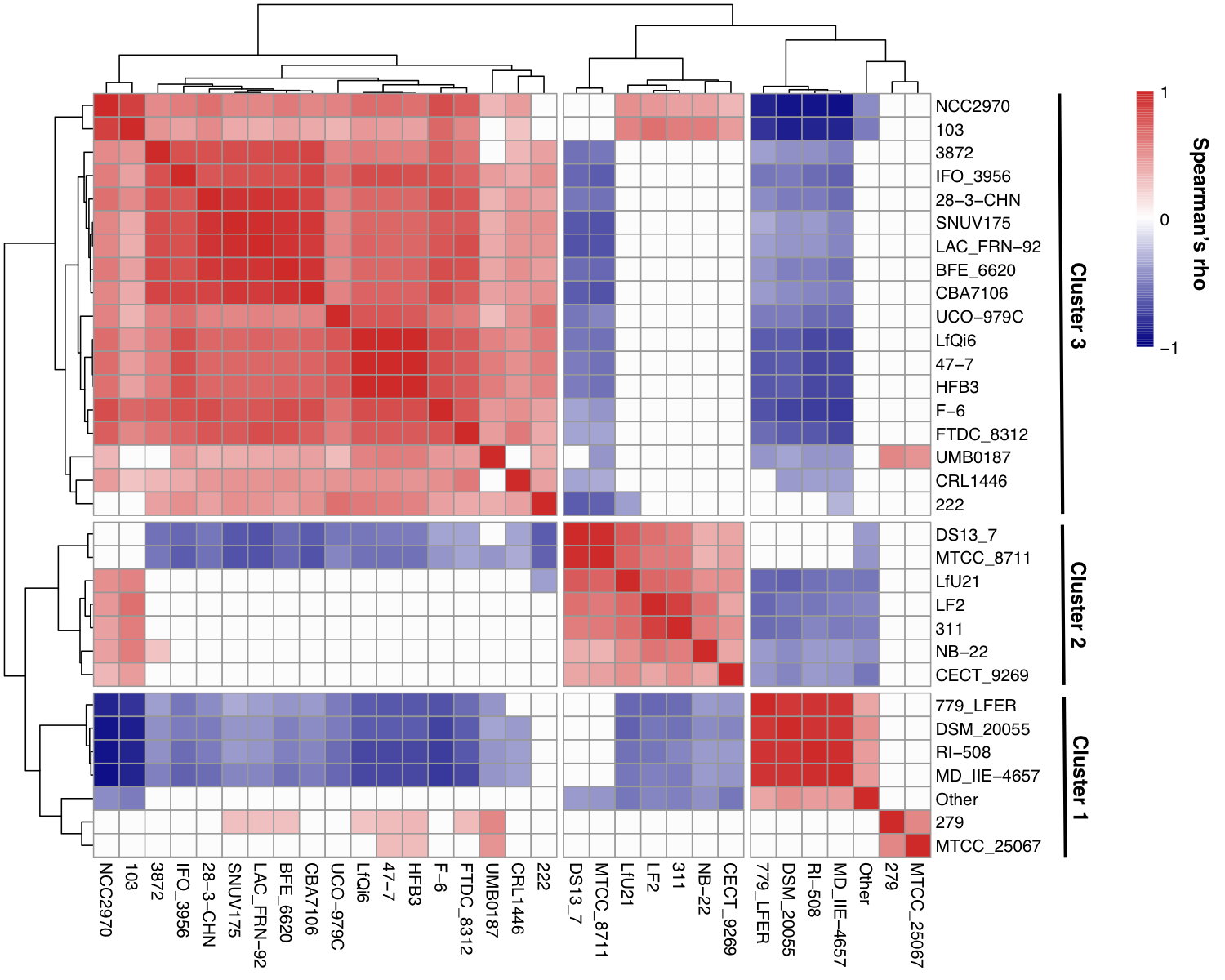

**Supplementary Figure 3. Strain clusters from *Lactobacillus fermentum* found in the bioethanol process.** Spearman’s correlation among the *L. fermentum* strains (FDR < 0.05) and the three strain clusters of *L. fermentum*. Hierarchical clustering among strains was performed on Euclidean distances. Strain abundance dissimilarity among different samples was calculated by Euclidean distances and hierarchical clustering was performed.
